# Using synthetic RNA to benchmark poly(A) length inference from direct RNA sequencing

**DOI:** 10.1101/2024.10.25.620206

**Authors:** Jessie J-Y Chang, Xuan Yang, Haotian Teng, Benjamin Reames, Vincent Corbin, Lachlan Coin

**Affiliations:** Department of Microbiology and Immunology, University of Melbourne at The Peter Doherty Institute for Infection and Immunity, Melbourne, VIC, 3000, Australia; Ray and Stephanie Lane Computational Biology Department, School of Computer Science, Carnegie Mellon University, Pittsburgh, Pennsylvania, United States of America, 15213; Department of Clinical Pathology, University of Melbourne, Melbourne, VIC, 3000, Australia

**Keywords:** Oxford Nanopore Technologies, poly(A) tail, estimation, segmentation, direct RNA sequencing

## Abstract

Polyadenylation is a dynamic process which is important in cellular physiology. Oxford Nanopore Technologies direct RNA-sequencing provides a strategy for sequencing the full-length RNA molecule and analysis of the transcriptome and epi-transcriptome. There are currently several tools available for poly(A) tail-length estimation, including well-established tools such as *tailfindr* and *nanopolish*, as well as two more recent deep learning models: *Dorado* and *BoostNano*. However, there has been limited benchmarking of the accuracy of these tools against gold-standard datasets. In this paper we evaluate four poly(A) estimation tools using synthetic RNA standards (Sequins), which have known poly(A) tail-lengths and provide a valuable approach to measuring the accuracy of poly(A) tail-length estimation. All four tools generate mean tail-length estimates which lie within 12% of the correct value. Overall, *Dorado* is recommended as the preferred approach due to its relatively fast run times, low coefficient of variation and ease of use with integration with base-calling.

## Background

Polyadenylation is a co-/post-transcriptional process in which a string of adenine nucleotides is added to the nascent messenger RNA molecule by enzymes such as polyadenylate (poly(A)) polymerases. This process is thought to increase the stability of the mRNA molecule [1], assist in export of the molecule from the cell nucleus [2] and is increasingly recognised as a dynamic process which influences timing and degree of protein production. As such, it is important to be able to measure polyadenylation accurately using a high-throughput assay to better understand its functional importance.

Oxford Nanopore Technologies (ONT) direct RNA-sequencing is an approach for single-molecule RNA-sequencing which does not require reverse transcription or polymerase chain reaction (PCR) amplification, thus avoiding amplification bias and retaining the original base and base-modification information [3-7]. Furthermore, full-length RNA molecules can be captured in one read, facilitating the identification of complex splicing patterns, RNA modifications and RNA secondary structures [8-13]. The nanopore sequencer records changes in ionic current as RNA passes through the pore in a custom FAST5/POD5 file. This raw data is then converted into sequence data using a custom deep learning model, such as *Dorado* or *Chiron* [14].

Nanopore sequencing of native RNA provides an attractive approach for measuring single-molecule poly(A) tail length. Thus, there have been several tools developed for estimating poly(A) tail length from raw nanopore signal (**Table 1**), including *nanopolish* [15], *tailfindr* [10], *Dorado* (developed by ONT) [16] as well as our in-house tool *BoostNano* (details described in **Supplementary Information**) [14]. There have been limited attempts to benchmark poly(A) tail length inference using gold-standard datasets with known poly(A) tail lengths.

**Table 1.** Description of each poly(A) tail estimation tool benchmarked in this study.

| Tool | Description | Reference |
| --- | --- | --- |
| *BoostNano* | Convolutional Neural Network (CNN)-Recurrent Neural Network (RNN)-Connection-ist Temporal Classification (CTC) architecture from *Chiron* basecaller used to find boundaries of poly(A) in raw signal | Teng et al., 2018. [14] |
| *tailfindr* | *R* tool, which uses the unaligned raw FAST5 data to estimate the poly(A) lengths via using the raw signal slope to refine the boundaries of potential poly(A) stretches and | Krause et al., 2019 [10] |
|  | normalization with the read-specific nucleotide translocation rate |  |
| *nanopolish* | Utilizes a predictive model in which a hidden Markov model (HMM - performs segmentation of the raw sequencing signal) and an estimator of the translocation rate are combined | Simpson et al., 2017 [15] |
| *Dorado* | Searches for the boundaries in the raw signal and estimates the poly(A) tail-length by taking into account the samples/base information, with adjustment for overestimation of the poly(A) tail. Primarily a basecalling tool, incorporates the poly(A) tail estimation during the basecalling itself | Oxford Nanopore Technologies [16] |

RNA Sequins are synthetic *in vitro* transcribed RNA [17]. RNA Sequins are transcribed from an artificial chromosome which comprises 78 gene loci split into two classes which have either a 30 bp (R1) or a 60 bp (R2) poly(A) tail. We have previously sequenced several direct RNA libraries which consisted of a mixture of host cell-line RNA together with spiked-in RNA sequins (BioProject: PRJNA675370) [12]. In this paper we re-use a subset of this data to benchmark poly(A) tail length estimation using *BoostNano, tailfindr, nanopolish* and *Dorado*.

### Performance evaluation between *BoostNano, tailfindr, nanopolish* and *Dorado*

To compare the estimation performance of *BoostNano, tailfindr* v1.4, *nanopolish* v0.13.3 and *Dorado* v0.5.3, we tested these tools on two Sequin testing sets with known poly(A) tail lengths: R1 set with 30 nucleotide (nt) tails and R2 set with 60 nt tails (**Figure 1 & Table 1**) [17]. The four methods display a similar pattern in the density distribution, with a prominent normal-like peak near the expected poly(A) length, but also with a over-representation of shorter poly(A) tails, ranging at approximately ∼0-10 nt (**Figure 1**). This peak was more prominent at ∼0-3 nt in *BoostNano*, whereas the early peaks for *tailfindr, nanopolish* and *Dorado* were positioned at ∼10 nt. We expected that these shorter peaks were derived from either fragmentation of the transcript, mis-priming of internal poly(A) stretches or degradation of the poly(A) tails. To test this, we inspected reads with <10 nt poly(A) tails and observed that the majority (68%) aligned within 20 nt of the 3’ end of the Sequin transcripts. (**Figures 2a & b**). This suggested that the majority of these shorter poly(A) tails is due to fragmentation/degradation of the poly(A) tail. However, the remaining 32% of reads showed truncations in the middle of the reference transcript (**Figure 2a**), consistent with mis-priming or fragmentation of the physical RNA. This was supported by stretches of adenine bases observed at the 3’ end of some of the truncated reads (**Figure 2c**). However, not all truncated reads showed poly(A) stretches (**Figure 2d**). Therefore, we hypothesize that some reads could have been able to be sequenced through mis-priming of these 3’ ends in regions with higher A-content.

**Figure 1.**
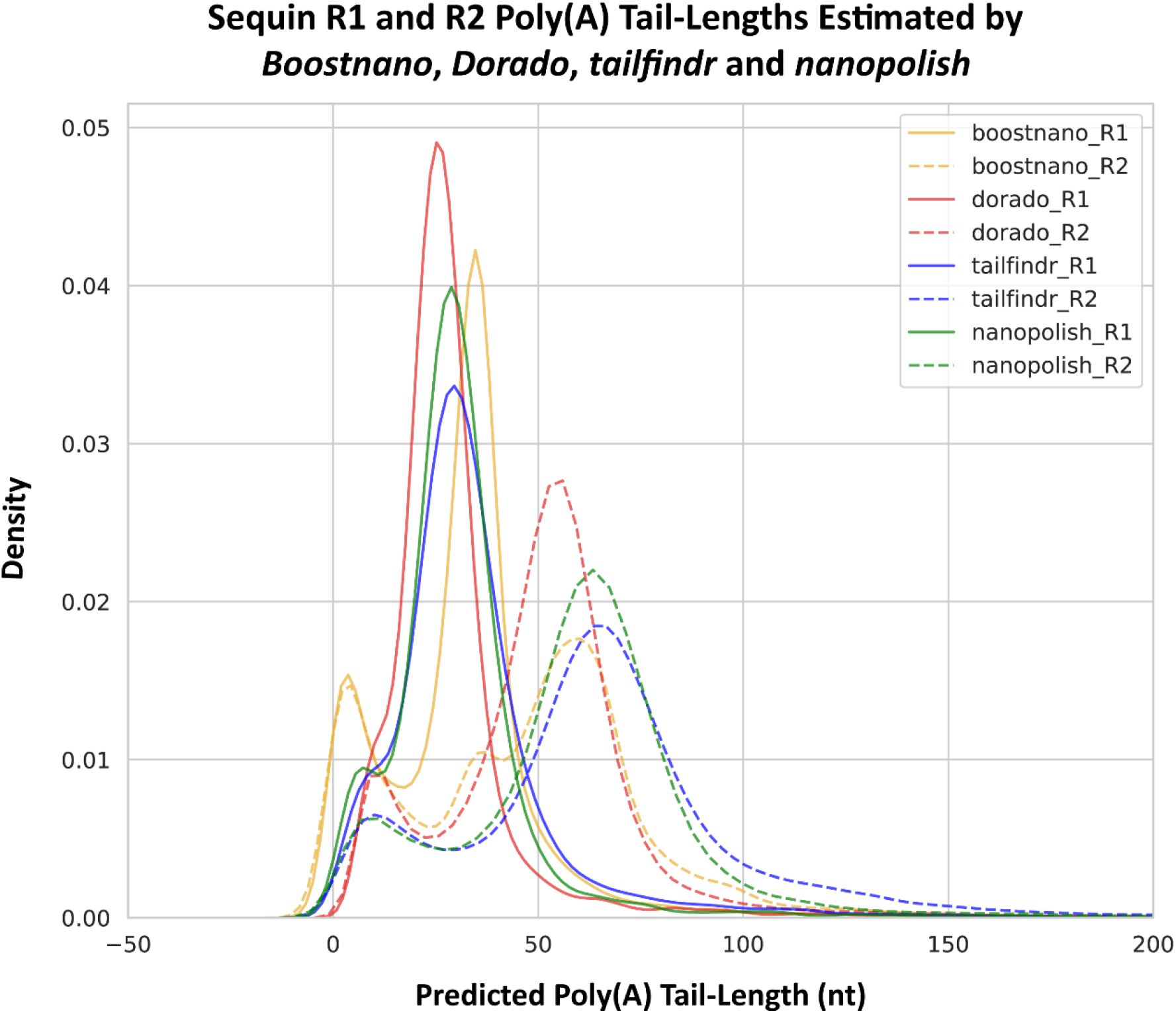
Density plots of estimated Sequin poly(A) tail-lengths. Outputs from *BoostNano* (yellow), *Dorado* (red), *tailfindr* (blue) and *nanopolish* (green). Expected poly(A) tail-lengths for R1 (solid lines) and R2 Sequins (dashed lines) are 30 nt and 60 nt, respectively. X-axis shows the predicted poly(A) tail-lengths of all reads and Y-axis reveals the density of the poly(A) tail-lengths.

**Figure 2.**
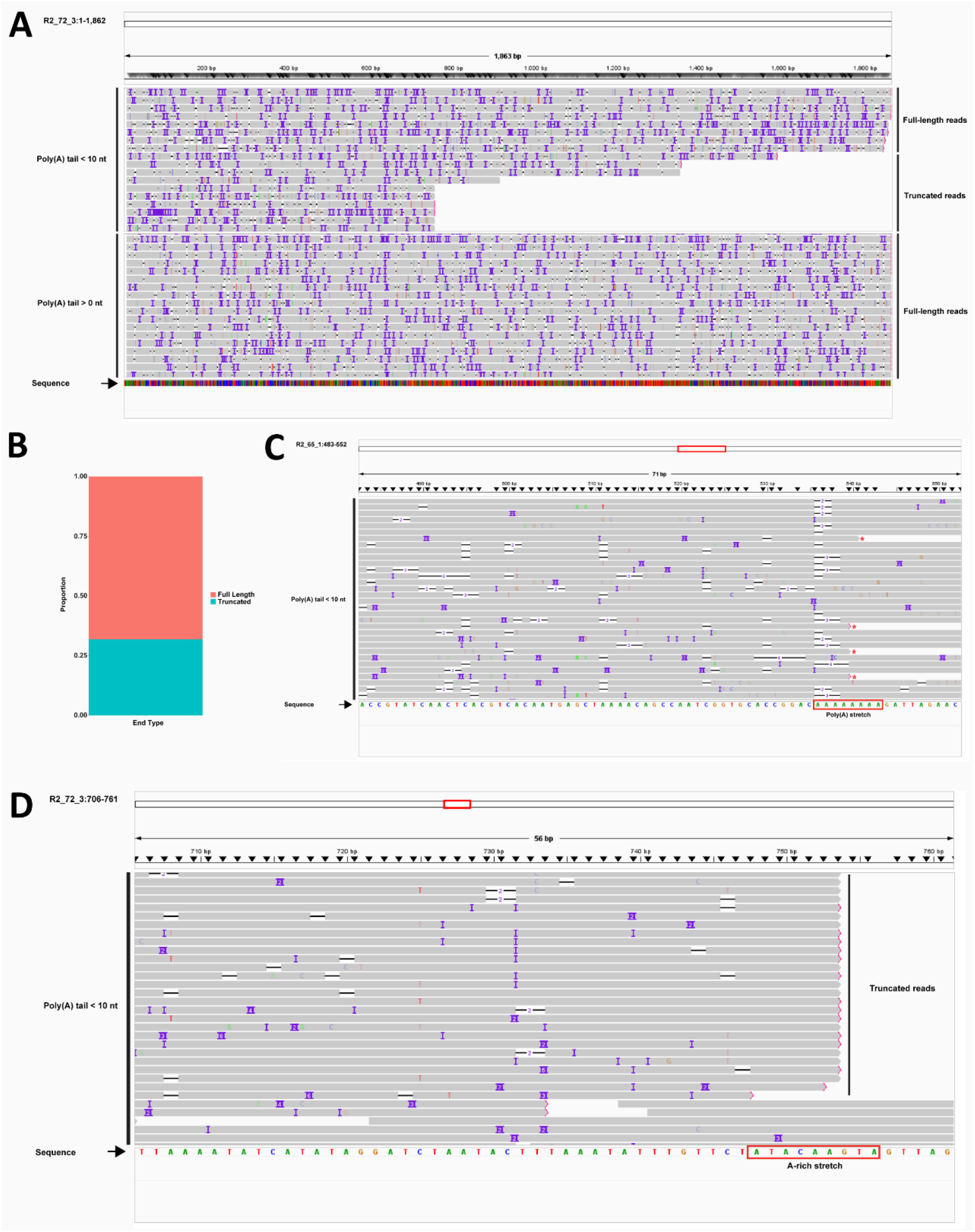
A representative subset of reads mapped to Sequins from the Vero 2 hpi dataset visualized with the Integrative Genomics Viewer (IGV) tool. **A**) A representative subset of reads mapping to the R2_72_3 Sequin transcript in Vero 2 hpi dataset. First subpanel shows a subset of reads with <10 nt poly(A) tails (estimated by *BoostNano*), showing that reads with <10 nt poly(A) tails are derived from both reads which have intact and also fragmented 3’ ends. The second panel shows a representative subset of full-length reads in the entire Vero 2 hpi dataset (i.e. poly(A) tails >0 nt), which have intact 3’ ends. **B)** Proportion of reads with truncated vs full-length 3’ ends in the Vero 2 hpi dataset with poly(A) tail-lengths <10 nt (estimated by *BoostNano*). **C)** Truncated reads with poly(A) tails <10 nt (estimated by *BoostNano*) and 3’ ends ending across an internal poly(A) stretch, mapped to R2_65_1 Sequin transcript marked with ^*^ (in red). **D)** The truncated reads with <10 nt poly(A) tails (estimated by *BoostNano*) which do not correlate to homopolymer poly(A) stretches at the 3’ end, mapped to the R2_72_3 Sequin transcript. Each grey line indicates a read. “Sequence” indicates the sequence of bases which form the transcript, where A = green, T = red, G = yellow and C = blue. The red box at the top of the figure indicates the zoomed out proportion of the reference transcript.

The explanations above partially explain the earlier peak (∼0-10 nt) in the density distribution (**Figure 1**) in all four methods, however, *BoostNano* particularly showed the mode of the peak presenting at even shorter poly(A) tail-lengths than *tailfindr, nanopolish* and *Dorado*. As the ONT reverse transcription adapter (RTA) used for reverse transcribing the native RNA strand has 10 polythymine (poly(T)) bases, it is likely that the minimum detection limit of poly(A) tails is 10 nt, which matches the ∼10 nt peak with *tailfindr, nanopolish* and *Dorado*. We expect that the poly(A) tails shorter than 10 nt occur due to potential truncation of the poly(T) stretch of the RTA. Interestingly, upon investigating these earlier peaks, we found that *Dorado* excludes reads which are retained in the analysis by *BoostNano*, despite them being classified as passed reads (**Figure 3**). While an earlier peak of <10 nt was the most prominent amongst reads discarded by *Dorado*, we also observed peaks of ∼40 nt and ∼60 nt (according to *BoostNano*) amongst reads discarded by *Dorado*. Therefore, *Dorado* appears to be a more conservative approach than *BoostNano*.

**Figure 3.**
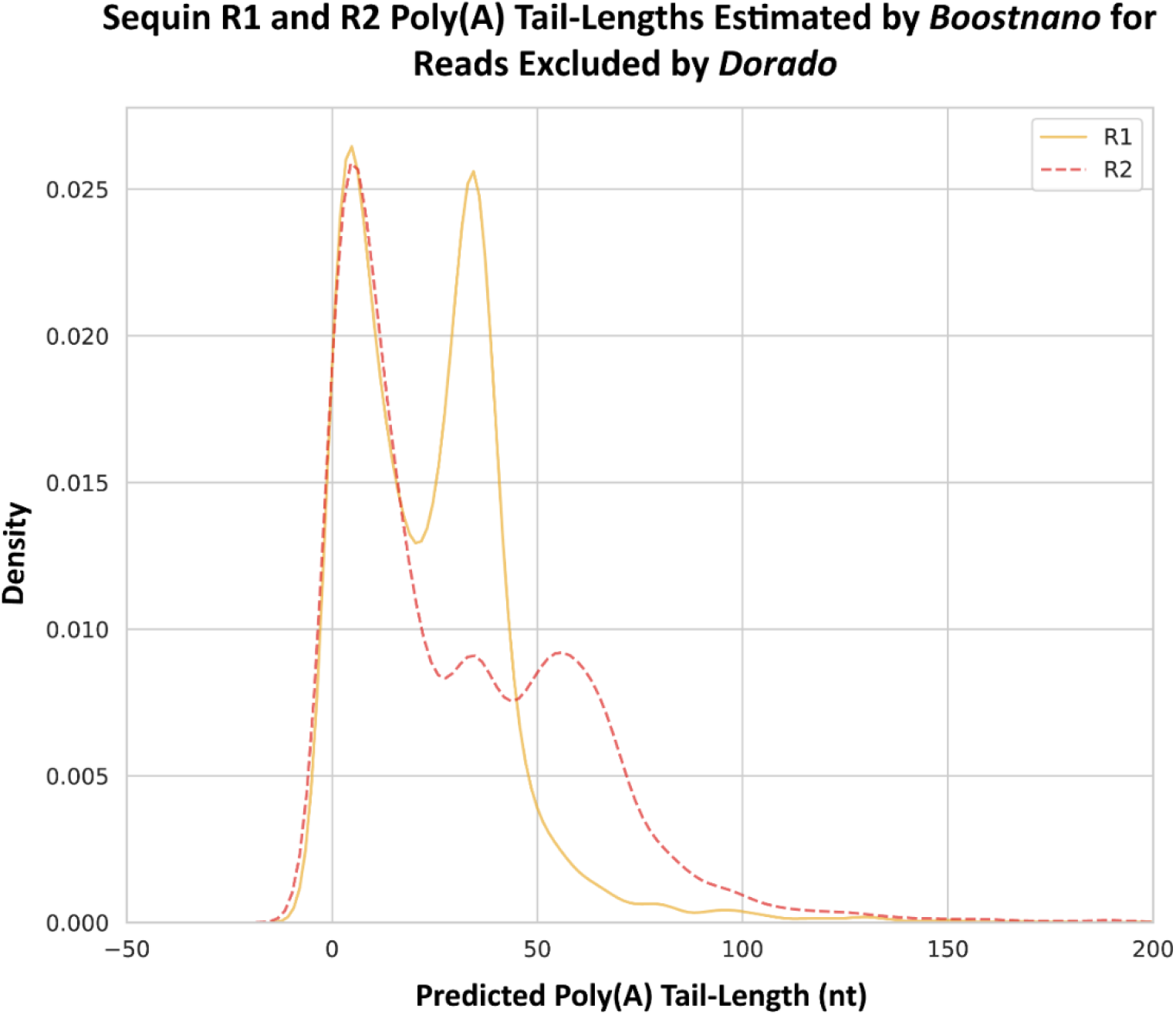
Reads which have been filtered out by *Dorado* but retained in the *BoostNano* output. R1 Sequin reads are indicated in yellow, and R2 Sequin reads are indicated in red. X-axis shows the predicted poly(A) tail length in nucleotides (nt), and the Y-axis shows the density distribution.

Following this, to estimate the accuracy of each method, we modelled the greatest peak as a normal distribution for each method and compared the mean and standard deviation (SD). The comparison was performed for both R1 and R2 Sequin sets (**Table 2**). In order to determine which normal distribution fit the peak best, we found the parameters (mean, SD) which minimize the root mean square error between the candidate normal distribution and the density distribution for an interval of 10 nt to the right of the mode. This is to remove biases due to an over-representation of the short-tailed reads. The peaks also lose their normal-like behavior for larger values. When comparing the results of the tools in both R1 and R2 testing sets, the deep learning tools - *BoostNano* and *Dorado* exhibited a more concentrated distribution around the mean, while *nanopolish* and *tailfindr* generally showed a wider spread of results (**Table 2**). With the R1 testing set, which had the shorter poly(A) tail-length, *tailfindr* showed the closest mean compared with the expected mean, and *BoostNano* showed the closest mean with the longer R2 testing set.

**Table 2.** Summary statistics for poly(A) tail-length estimation by four tools (*BoostNano, Dorado, tailfindr* and *nanopolish*) on two Sequin testing sets (R1 and R2).

| Testing set | Method | Read Count | Mean (nt) | SD (nt) |
| --- | --- | --- | --- | --- |
| R1 (30 nt) | *BoostNano* | 14130 | 34.9 | 4.3 |
|  | *Dorado* | 9536 | 26.1 | 5.7 |
|  | *tailfindr* | 14106 | 29.3 | 8.8 |
|  | *nanopolish* | 9036 | 28.2 | 7.8 |
| R2 (60 nt) | *BoostNano* | 42196 | 59.3 | 11.3 |
|  | *Dorado* | 29568 | 54.5 | 9.5 |
|  | *tailfindr* | 41173 | 65.4 | 12.6 |
|  | *nanopolish* | 23645 | 64.1 | 11.0 |

Next, we compared the computational time required by each method to predict the tail-length of 4000 reads. For *BoostNano* and *Dorado*, we used one graphics processing unit (GPU) with 16G allocated RAM, while for *nanopolish* and *tailfindr*, which doesn’t have the option to be run on GPU, we used one central processing unit (CPU) with 16G RAM and 1 thread for *nanopolish. Dorado* and *nanopolish* showed rapid computational times under 3 minutes (m), whereas *BoostNano* revealed the longest computational time at ∼16 m and 52 seconds (s) (**Table 3**). *BoostNano* also offers the option of using the Application Programming Interface (API) call instead of the direct method, which omits the file copy implemented in the direct approach, reducing the run time to 8 m 8 s.

**Table 3.** Computational time efficiency to process 4000 reads with 1 GPU/CPU.

| Method | Execution time (4000 reads) | GPU/CPU |
| --- | --- | --- |
| *BoostNano (direct)* | 16 m 52 s | GPU |
| *BoostNano (API call)* | 8 m 8 s | GPU |
| *Dorado* | 0 m 51 s | GPU |
| *tailfindr* | 10 m 56 s | CPU |
| *nanopolish* | 2 m 5 s | CPU |

## Discussion

In this technical note, we assessed the predictive performance of 4 poly(A) tail-length estimation tools - *tailfindr, nanopolish, BoostNano* and *Dorado* on two separate testing sets with known poly(A) tail-lengths. *BoostNano* and *tailfindr* tools provided estimation of the starting and ending positions of the poly(A) tails in event space while this information was absent in *Dorado* outputs. All four methods estimated the length of the tails within 1.2 SD as calculated above (**Table 2**). However, there were differences in the accuracy of the methods. On the R1 dataset, *BoostNano* showed a tighter distribution with the smallest SD, but its peak was the furthest from the correct value. *tailfindr* had the most accurate estimation but also the largest error interval. On the R2 dataset, however, the *BoostNano* estimation was the most accurate, with a fairly large SD, while *Dorado, nanopolish* and *tailfindr* were equally inaccurate with *Dorado* having a slightly smaller SD. Furthermore, *Boostnano* is more lenient in keeping reads for poly(A) estimation than *Dorado*. Overall, our results suggest that the four tools investigated in this study - *BoostNano, tailfindr, nanopolish* and *Dorado* have similar performance with their accuracy varying from one dataset to the other, with a potential length bias. The only obvious difference is in the speed of execution, where *Dorado* and *nanopolish* surpasses the rate of *BoostNano* or *tailfindr* (**Table 3**). Therefore, we expect *Dorado* to be implemented as the default method of poly(A) tail estimation in the near future, with the rapid estimation timeframe, comparable estimation lengths to other tools, conservative nature and the added benefit of ease of obtaining this information during basecalling.

This work demonstrates the value of having access to synthetic RNA molecules with known poly(A) tail-lengths for validating the accuracy of poly(A) tail estimation algorithms. As methods improve, we anticipate that these datasets will be valuable for assessing improvements in estimation of poly(A) tails.

## Methods

### Dataset

This study utilized publicly available ONT dRNA-seq datasets involving SARS-CoV-2-infected continuous cell lines (Vero, Calu-3 and Caco-2) derived from our previous study, with synthetic RNA - Sequins (BioProject: PRJNA675370) [12]. Briefly, Vero (African green monkey kidney epithelia), Calu-3 (Human lung adenocarcinoma epithelia) and Caco-2 (Human colorectal adenocarcinoma epithelia) cells were cultured in 6-well tissue culture plates at 37°C, 5% (v/v) CO_2_. The Australian ancestral strain of SARS-CoV-2 (SARS-CoV-2/human/AUS/VIC01/2020) was used to infect these cells at a multiplicity of infection (MOI) of 0.1 and the cells were harvested at 0, 2, 24 and 48 hours post-infection (hpi). The total RNA was extracted and 6 μg of total RNA for Vero cells and 3 μg of total RNA for Calu-3 and Caco-2 cells + 10% of expected mRNA (5% of total RNA) of Sequins [17] were added to the sample pool. The RNA was sequenced using the ONT Direct RNA Kit (SQK-RNA002), on R9.4.1 flow cells via the ONT MinION/GridION. For the purposes of this study, infected datasets from 24 and 48 hpi from all three cell lines and additionally 2 hpi from Vero cells were analyzed.

## Findings

## Availability of supporting source code and requirements

Project name: BoostNano

Project home page: https://github.com/haotianteng/BoostNano

Operating system(s): Platform independent

- Programming language: Python
- Other requirements: Pytorch
- License: Mozilla Public License, v. 2.0

## Data Availability

The data sets supporting the results of this article are available in the NCBI repository, BioProject: PRJNA675370.

## Declarations

## List of abbreviations

Poly(A): polyadenylate
ONT: Oxford Nanopore Technology
PCR: Polymerase Chain Reaction
HMM: Hidden Markov Model
CNN: Convolutional Neural Network
RNN: Recurrent Neural Network
CTC: Connection-ist Temporal Classification
SD: Standard deviation
m: minutes
s: seconds

## Ethics approval and consent to participate

Not applicable.

## Consent for publication

Not applicable.

## Authors’ contributions

Conceptualization –J.J.-Y.C., H.T., V.C., L.C.

Methodology – H.T., V.C., L.C.

Software – H.T., V.C., L.C.

Validation - H.T., X.Y., V.C., L.C.

Formal analysis - H.T., B.R., X.Y., V.C., L.C.

Investigation - H.T., B.R., X.Y., V.C., L.C.

Resources - J.J.-Y.C., H.T., V.C., L.C.

Data Curation - J.J.-Y.C., V.C., L.C.

Writing - Original Draft - J.J.-Y.C., H.T., X.Y., V.C., L.C.

Writing - Review & Editing - J.J.-Y.C., H.T., X.Y., V.C., L.C.

Visualization – J.J.-Y.C., H.T., X.Y., V.C., L.C.

Supervision – V.C., L.C.

Project administration – J.J.-Y.C., V.C., L.C.

Funding acquisition – V.C., L.C.

## Acknowledgements

This research was supported by The University of Melbourne’s Research Computing Services and the Petascale Campus Initiative.

## Authors’ information

Not applicable.

