## Supplementary Figures and Tables for "Using synthetic RNA to benchmark poly(A) length inference from direct RNA sequencing"

**Supplementary Information**

### Model structure of *Boostnano*

*BoostNano* is a machine learning-based tool based on *Chiron* - an end-to-end nanopore sequencing basecaller established using a deep learning CNN+RNN+CTC structure [13], which preprocesses ONT raw reads in FAST5 files before basecalling. Unlike traditional methods that involve basecalling, FASTQ formatting, and sequence alignment, *BoostNano* directly extracts, transforms, and analyzes the raw electrical current signal information within FAST5 files in its neural network model. The process involves segmenting the adaptor, poly(A) stalls, as well as transcription from the raw signal [13]. The inner detection model of *BoostNano* consists of two branches, a main branch and a side branch, each with three conventional layers and corresponding activation functions. The signal is processed in a streaming manner. Current signal block is fed into the side branch, and the output is then concatenated with the previous hidden state processed by the main branch, predicting the current signal's state (adapter, poly(A) stalls, or transcription). This approach significantly reduces the detection time and can process approximately 200 to 400 direct RNA reads per minute per GPU. An output example of the segmentation plot for one RNA read generated by *BoostNano* is shown in **Figure S1**. The three red-line stalls indicate the locations of the adapter, poly(A) tail starting site, and poly(A) tail ending site.

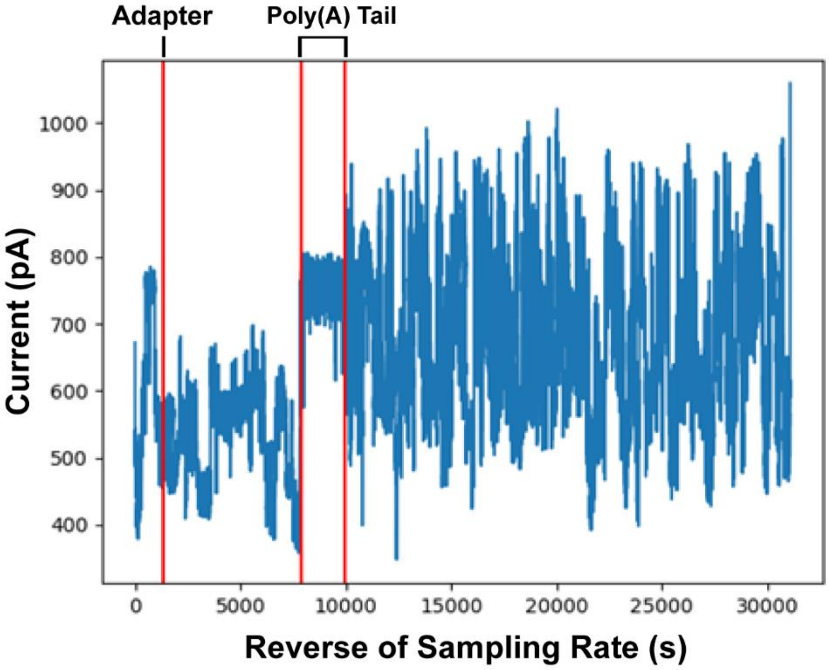


**Figure S1. An output example of the segmentation plot using *BoostNano*.** The first red line indicates the start of the adapter, second line indicates the position of the start of the poly(A) tail and the third line indicates the position of the end of the poly(A) tail. The Y-axis is the electric current (picoAmperes (pA)) and the X-axis is the reverse of sampling rate (1/4000 seconds (s)), as Nanopore sequencing samples at 4000 events per second.

The general model structure of *BoostNano* is as follows (**Figure S2**):

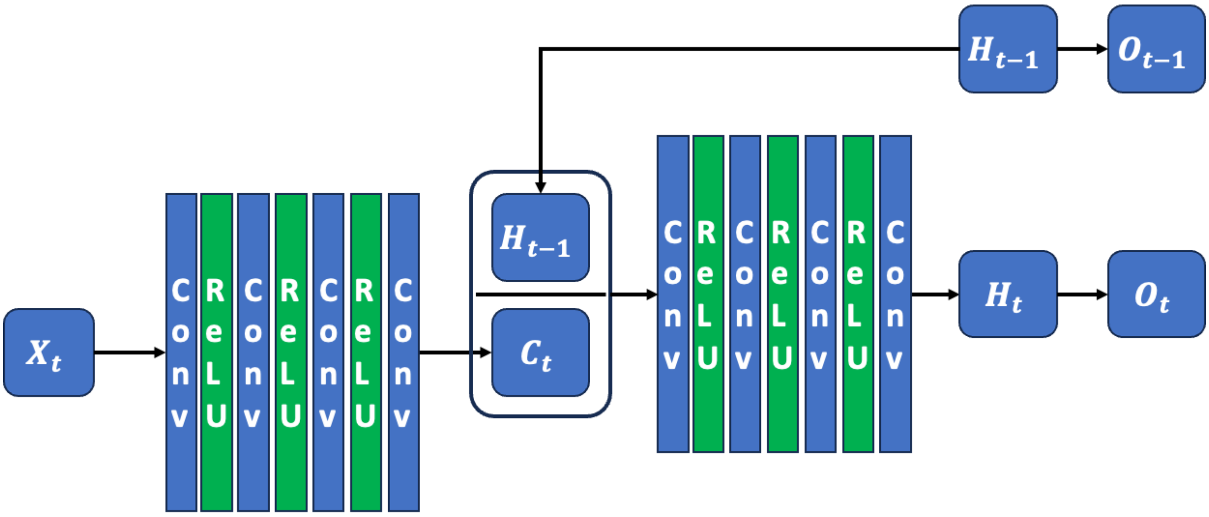


**Figure S2. General model structure of *BoostNano*.**

Given the signal $X = \{x_t; t = 1, \dots, T\}$, *BoostNano* predicts the state of the signal at time $t$. Specifically, the state $\mathbf{h}_t = \{A1, A2, P, T\}$ is determined, where $A1$ denotes the sequencing adapter, $A2$ is the adapter added during poly(A) tailing, $P$ represents the poly(A) tail, and $T$ is the targeted transcription signal. The *BoostNano* model employs a continuous convolutional architecture. At each step, the neural network combines the previous hidden states with the current signal segments to predict the signal's state, described as $\mathbf{h}_t = \mathbf{g}_1(\mathbf{f}(x_{t:t+k}) \mid \mathbf{h}_{t-1})$ for $t > 1$. Otherwise, for the initial step, it is $h_0 = g_2\big(f(x_{0:k})\big)$, where $k$ represents the width of the receptive field, and $\mathbf{f}, \mathbf{g}_1, \mathbf{g}_2$ are three distinct convolutional neural networks.

### Model training

The training data was labeled by human agents using a labeling GUI developed by us. During the labeling phase, each agent was required to identify the time points where transitions occurred, such as from the sequencing adapter to the poly(A) tailing adapter, then to the poly(A)-tail, and finally to the transcription RNA signal. To minimize personal bias, the data were labeled by three different human agents. The number of reads labeled by each agent was 436, 1021, and 1270 respectively. In total, 2186 reads out of 2727 passed our data review and were included in the final training dataset. Additionally, 1165 reads were labeled and designated as the validation dataset. The models were trained using the Adam optimizer and trained for two epochs.

### Model inference

During inference, the model outputs the probability of the signal state at each time point, and the maximum likelihood under the state Markov chain ($s \to A1 \to A2 \to P \to T$) is calculated to obtain the final signal segmentation.
